# Depletion of *ALMS1* affects TGF-β signalling pathway and downstream processes such as cell migration and adhesion capacity

**DOI:** 10.1101/2021.12.19.473196

**Authors:** Brais Bea-Mascato, Elena Neira-Goyanes, Antía Iglesias-Rodríguez, Diana Valverde

## Abstract

**Background:** *ALMS1* is a ubiquitous gene associated with Alström syndrome (ALMS). The main symptoms of ALMS affect multiple organs and tissues, generating at last, multi-organic fibrosis in the lungs, kidneys and liver. TGF-β is one of the main pathways implicated in fibrosis, controlling the cell cycle, apoptosis, cell migration, cell adhesion and epithelial-mesenchymal transition (EMT). Nevertheless, the role of *ALMS1* gene in fibrosis generation and other implicated processes such as cell migration or cell adhesion via the TGF-β pathway has not been elucidated yet.

**Methods:** Initially, we evaluated how depletion of *ALMS1* affects different processes like apoptosis, cell cycle and mitochondrial activity in HeLa cells. Then, we performed proteomic profiling with TGF-β stimuli in HeLa ALMS1 −/− cells and validated the results by examining different EMT biomarkers using qPCR. The expression of these EMT biomarkers were also studied in hTERT-BJ-5ta ALMS1 −/−. Finally, we also evaluated the SMAD3 and SMAD2 phosphorylation and cell migration capacity in both models.

**Results:** Depletion of *ALMS1* generated apoptosis resistance to thapsigargin (THAP) and C2-Ceramide (C2-C), and G2/M cell cycle arrest in HeLa cells. For mitochondrial activity, results did not show significant differences between *ALMS1 +/+* and *ALMS1 −/−.* Proteomic results showed inhibition of downstream pathways regulated by TGF-β. The protein-coding genes (PCG) were associated with processes like focal adhesion or cell-substrate adherens junction in HeLa. *SNAI1* showed an opposite pattern to what would be expected when activating the EMT in HeLa and BJ-5ta. Finally, in BJ-5ta model a reduced activation of SMAD3 but not SMAD2 were also observed. In HeLa models no alterations in the canonical TGF-β pathway were observed but both cell lines showed a reduction in migration capacity.

**Conclusion:** *ALMS1* has a role in controlling the cell cycle and the apoptosis processes. Moreover, the depletion of *ALMS1* affects the signal transduction through the TGF-β and other processes like the cell migration and adhesion capacity.

## INTRODUCTION

*ALMS1* is a ubiquitous gene with 23 exons located on chromosome 2p13. Mutations in this gene are related to Alström syndrome (ALMS; OMIM #203800), a rare, autosomal and recessive disease with a prevalence of 1-9 cases per 1.000.000 inhabitants and approximately 950 cases reported worldwide (Orphanet). The main symptoms of ALMS affect multiple organs and tissues [1], such as cone-rod retinal dystrophy, dilated cardiomyopathy (DCM) and hypertension [2], hypertriglyceridemia, type II diabetes mellitus (T2D) [3], and hepatic, renal and pulmonary dysfunction caused by multi-organic fibrosis [1], [4], which is the main cause of death in these patients.

The protein encoded by *ALMS1* plays a structural role in the eukaryotic cell centrosomes [5], a microtubule (MT)-nucleating organelle strongly related to primary cilia. The ALMS1 basal body location [5], [6] has been related to different functions controlled by the primary cilium such as centrosome cohesion, cell cycle control and endosomal trafficking [6]–[9].

Although numerous roles of ALMS1 have been described at both ciliary and extraciliary levels, the regulation of the pathways underlying these cellular processes remain unknown. Recently, our group has established the relationship between the *ALMS1* gene and the TGF-β signaling pathway [10], which could be related to the development of fibrosis in these patients [11], [12]. TGF-β is a signalling pathway regulated by clathrin-dependent endocytosis (CDE) in the pocket region of primary cilium [13], and coordinates cell cycle, apoptosis and migration processes such as epithelium-mesenchyme transition (EMT) [14]. This pathway acts in close coordination with other signalling cascades such as Wnt, Hippo, Notch, Hedgehog (Hh), mitogen-activated protein kinase (MAPK), and phosphoinositide 3-kinase (PI3K)-Akt, resulting in many cases in complex regulatory networks dependent on the primary cilium [15].

Stimulation of the TGF-β pathway can activate two main cascades: the canonical (SMAD-dependent) pathway and the non-canonical (SMAD-independent) pathway [16]. The canonical pathway is coordinated by transcription factors called R-SMAD (SMAD2 and SMAD3) which join co-SMAD (SMAD4) to activate the expression of several genes such as *SNAI1* [17], [18], a key modulator of EMT [19], [20]. This pathway is inhibited by an I-SMAD, SMAD7, which inhibits R-SMAD signalling in a variety of ways, binding to TGF-β receptors or directly in the nucleus, avoiding the binding of the R-Smad-Smad4 complex to DNA [21]. On the other hand, the non-canonical activation pathway includes a wide group of pathways such as p38 MAPK, PI3K/AKT, or Rho GTPases [16]. Recently, studies have linked the abolition of several centrosomal proteins such as ALMS1 and CEP128 to the inhibition of the SMAD-dependent canonical pathway in zebrafish and various human cell types such as hTERT-RPE1 and ciliated human foreskin fibroblasts (hFF) [10], [22]. However, deciphering how centrosomal proteins coordinate and regulate the TGF-β signalling pathway and their downstream processes is a challenge.

Understanding how the absence of *ALMS1* influence cell migration and adhesion processes such as EMT through the TGF-β pathway is key to understanding how the multi-organ fibrosis occurs in these patients.

In this article, we analysed how the lack of *ALMS1* could affect different cell processes such as mitochondrial activity, cell cycle, apoptosis, and signal transduction through the TGF-β pathway. Moreover, we evaluated the expression pattern for different EMT biomarkers after TGF-β pathway stimulation in two independent CRISPR-KO cell models (HeLa and BJ-5TA) for *ALMS1.*Finally, we measured p-SMAD3 and p-SMAD2 activation and migration capacity in in the presence and absence of the TGF-β1 ligand.

## MATERIALS AND METHODS

### Cell culture

HeLa human cervical cancer cell line from American Type Culture Collection (ATCC), hTERT BJ-5ta human dermal fibroblasts (ATCC) and 293T human embryonic kidney cell line from ATCC were used in this study. HeLa and 293T cells were maintained in Dulbecco’s minimum essential medium (DMEM, Gibco, Invitrogen, NY, USA), supplemented with 10% fetal bovine serum (FBS) (Gibco, Invitrogen, NY, USA) and 2% penicillin/streptomycin (P/S) (Gibco, Invitrogen, NY, USA). In the case of BJ-5ta line, cells were maintained in DMEM:199, supplemented with 10% of FBS, 2% P/S and Hygromycin B (0.01mg/mL). All cell lines were grown at 37°C with 5% CO_2_ for up to one month. After this, the cells were discarded, and a new culture was started.

### Crispr assay and edition validation

HeLa cells were edited with a dual episomal system for homology directed repair (HDR) (Santa Cruz Biotechnology, Inc., Dallas, USA). Cells were transfected with Lipofectamine 2000 (Thermo Fisher, Waltham, EE. UU) with 3 gRNAs directed to *ALMS1* exons 1 and 3 (Supplementary Table S1) following the manufacture’s protocol. After plasmid transfection, cells were selected with puromycin (1 μg/mL) for 5 days.

**Supplementary Table S1.**
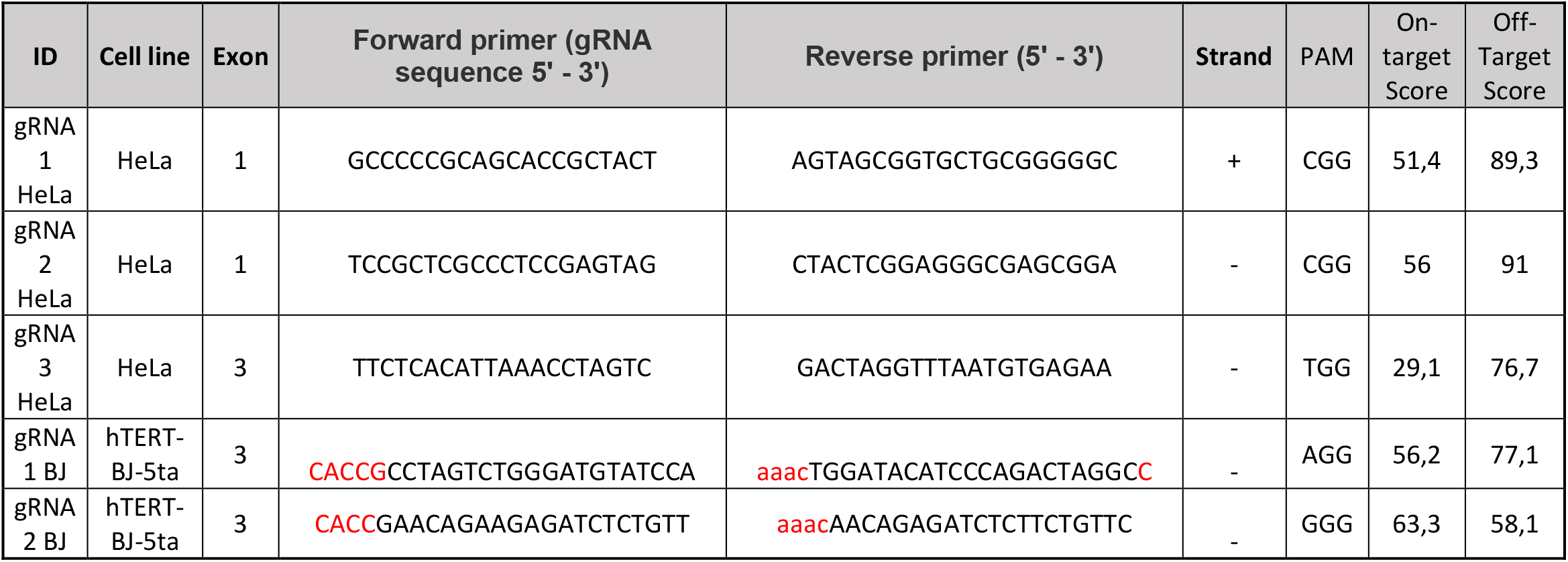
gRNAs of the constructs used for gene editing. The forward and reverse sequences are shown in the table. The red colour represents the base pairs added for ligation in the plasmid.

BJ-5ta cells were edited using a lentivirus system according to the protocol described by Sanjana et al [23], with some modifications. Check supplemental material for more information. After lentiviral transduction, cells were selected with puromycin (1μg/mL) for 5 days.

Surviving cells were expanded and isolated to obtain individual clones. Of the isolation was performed using Fluorescence-Activated Cell Sorting (FACS) with Hoechst (BD Bioscience, San José, USA) in the flow cytometer FACS ARIA III (BD Bioscience, San José, USA). Inhibition of gene expression was validated by qPCR in a StepOnePlus instrument (Thermo Fisher, Waltham, USA). A commercial probe TaqMan^®^ Gene Expression Assays (Hs00367316_m1) and PrimeTime^®^ qPCR Primer Assays (Hs.PT.56a.488948) were used to detect *ALMS1* in the HeLa and BJ-5ta models, respectively.

### Mitochondrial activity assay

Mitochondrial activity was evaluated using Presto Blue reagent (Thermo Fisher, Waltham, USA). Wild type (*ALMS1* +/+) and Hela Knockout cells (*ALMS1* −/−) were seeded in 96-well plates by triplicate in concentrations from 5 × 10^4^ to 1.5 × 10^3^ cells per well. The plates were incubated for 24 hours as previously described. After incubation, 10 μL of Presto Blue were added to each well and cells were incubated for 2 hours. Next, the plate was centrifuged at 1500 rpm for 5 min and the supernatant was transferred to a fluorescence plate (Corning, NY, EE. UU). Finally, the plates were measured in the Envision plate reader (Perkin-Elmer, Waltham, USA). This experiment was repeated 3 times on different days.

### Apoptosis assay

The apoptosis assay was performed in flat-bottom 96-well plates. HeLa *ALMS1* +/+ and *ALMS1* −/− cells were seeded in triplicate at a concentration of 3 × 10^4^ cells/per well and incubated 24 hours in DMEM 2%P/S and 2%FBS before treatment. Next, apoptotic stimuli were added at final concentration of 100 nM for thapsigargin (THAP) and 25μM for C2-ceramide (C2-C) in plates, followed by incubation at 37ºC for 24-48 hours respectively. After that, cells were washed twice with 100 μL PBS. 50 μL trypsin was added per well and the plate was incubated for 5 min at 37°C. After that, trypsin was inactivated with 50 μL DMEM 10% FBS 2% P/S, and cells were transferred to 96-well plates with U bottom. The plate was centrifugated at 1500 rpm for 5 min, the supernatant was discarded, and the cells were resuspended in 200 μL biding buffer with 2μL 7-AAD and 2μL annexin V-FITC (Biolegend Inc, San Diego, CA, USA). The plate was incubated for 15 min in darkness and measured in the flow cytometer FC 500 (Beckman-Coulter, Brea, CA, USA). This experiment was repeated 3 times on different days.

### Cell cycle assay

To determine cell cycle alterations, we studied sub-populations in the HeLa *ALMS1* −/− model with Hoechst staining (BD Bioscience, San José, USA). *ALMS1* +/+ and *ALMS1* −/− cells were seeded in a 6-well plate at a concentration of 3 × 10^5^ cells/per well and incubated overnight. The next day, 1μL Hoechst staining was added to the medium and the plate was incubated for 1 hour. Then, the medium was removed, cells were washed twice with 1 mL PBS per well, and trypsinised following the protocol described above. Cells were resuspended in PBS and analysed in the flow cytometer FACS ARIA III.

### Protein extraction for proteomic profiling

HeLa cells were seeded in 6-well plates by triplicate in DMEM 2% P/S, 2%FBS and incubated overnight for serum starvation. One the next day, cells were stimulated with rhTGF-β1 (2ng/mL; R&D Systems; 240-B) for 24 hours. After that, DMEM was removed, and wells were washed twice with PBS. Cells were scrapped in PBS. Then, cells were pelleted at 11000 rpm for 5 min, and 100 μL of RIPA Buffer were added to each sample. Samples were incubated on ice for 20 min in constant agitation. Finally, cell debris was pelleted by centrifugation for 30 min at 12000 rpm and 4 °C. Samples were aliquoted and stored at −80 °C until the next protocol’s step. Protein quantification was performed by Bradford microplate assay (Bio-Rad, Hercules, USA).

### Protein concentration, reduction, and alkylation

Initially, 20 μg of each sample was concentrated and stained with Coomassie in a 12% SDS-PAGE gel. Bands were cut and faded. Band distinction was performed by adding 100 μL of ammonium bicarbonate 25mM/50% acetonitrile and incubated 20 min. This step was repeated 4 times. After distinction, the supernatant was removed and 100 μL of ammonium bicarbonate 25mM-50% EtOH was added to samples, and incubated for 20 min. Then, the supernatant was removed and 100 μL EtOH was added and incubated for 15 min. In the next step, 200 μL 10mM DTT in ammonium bicarbonate were added and samples were incubated for 1 hour at 56°C. Then, DTT solution was removed and 200 μL iodoacetamide (IAA) 55mM in 50 mM ammonium bicarbonate was added. Samples were incubated again for 30 min at room temperature. Finally, the supernatant was removed, 200 μL ammonium bicarbonate 50mM were added and samples were incubated for 15 min.

### Protein digestion

After reduction and alkylation, samples were digested. 100 μL ammonium bicarbonate 25mM-50% acetonitrile (ACN) was added to each sample and incubated for 30 min. The supernatant was removed and 100 μL ACN 100%was added and incubated for 10 min. The supernatant was removed again and 15 μL Trypsin in ammonium bicarbonate 25mM were added to each sample. Samples were maintained at 4°C for 30 min. Finally, trypsin was removed, 5 μL ammonium bicarbonate 25 mM were added, and samples were incubated overnight.

### Peptide extraction

40 μL of trifluoroacetic acid (TFA) 0.5% were added to each sample and incubated for 30 min at room temperature. Then, we added 20 μL ACN 100% in each 1.5 mL tube and incubated for 15 min at 37°C. Finally, the samples were dried in a speed-vac at 45°C for 1 hour and stored at −20°C until the analysis.

### Mass Spectrometry and bioinformatic analysis

Samples were resuspended in 24 μL formic acid 0.1%. Then, 10 μL were injected into the HPLC (LC-ESI-MS/MS workflow using a C18 precolumn). Samples were analysed in a mass spectrophotometry Orbitrap ELITE (Thermo-Fisher, Waltham, USA) in long chromatography mode (4 hours). After that, protein identification was performed in the database UniProtKB/TrEMBL for humans, using the Proteome Discoverer software. Downstream analysis was performed with the R package Differential Enrichment analysis of Proteomics data (DEP package) [24]. Finally, Go Terms analysis was performed with Clusterprofiler package [25], and results were visualized with GOplot package [26]. All the code for the analysis was deposited in the GitHub repository (https://github.com/BreisOne/CRISPR_Hela_KO_Alms1).

The mass spectrometry proteomics data have been deposited in the ProteomeXchange Consortium via the PRIDE [27] partner repository with the dataset identifier PXD024964.

### RNA extraction and RT-PCR

HeLa cells were seeded in 6-well plates at a concentration of 2×10^5^ by triplicate in DMEM 2%P/S and 2%FBS and were incubated overnight for serum starvation. The next day, cells were stimulated with rhTGF-β1 (2 ng/mL) for 24, 48 and 72 hours. DMEM was removed and wells were washed twice with PBS. Cells were scraped in PBS and collected in a 1.5 mL tubes. For the RNA extraction, the NYZ total RNA isolation kit (NYZtech, Lisboa, Portugal) was used following the manufacture’s protocol.

After RNA elution, the sample concentrations were measured with Nanodrop (Thermo Fisher, Waltham, USA). Next, 300 ng of RNA were retrotranscribed to cDNA using the NZY First-Strand cDNA Synthesis kit (NYZtech, Lisboa, Portugal) following the manufacture’s protocol.

For the experiment with the BJ-5ta line, the same protocol was followed but using DMEM:199 and stimulating for 24 and 48 hours only.

### qPCR expression analysis

Samples were prepared in 96-well plates (Thermo Fisher, Waltham, EE. UU) adding 1μL of each primer, 12.5μL of PowerUp SYBR Green Master Mix (Thermo Fisher, Waltham, EE. UU), 1μL of cDNA and 4.5μL of H_2_0_d_ in a final volume of 20 μL per well. qPCRs were performed in StepOnePlus instrument using the following, cycling conditions: 10 min at 95°C, 40 cycles of 15 s at 95°C and 1 min at 60°C.Primers were synthesized on demand (IDT) (Supplementary Table S2). Samples were analysed by technical triplicates for each sample and gene. Cts for technical triplicates were averaged using the geometric mean and ΔΔCT method was applied for the normalization, using 2 housekeeping (HK) genes (*ALAS1* and *YWHAZ*).

**Supplementary Table S2.**
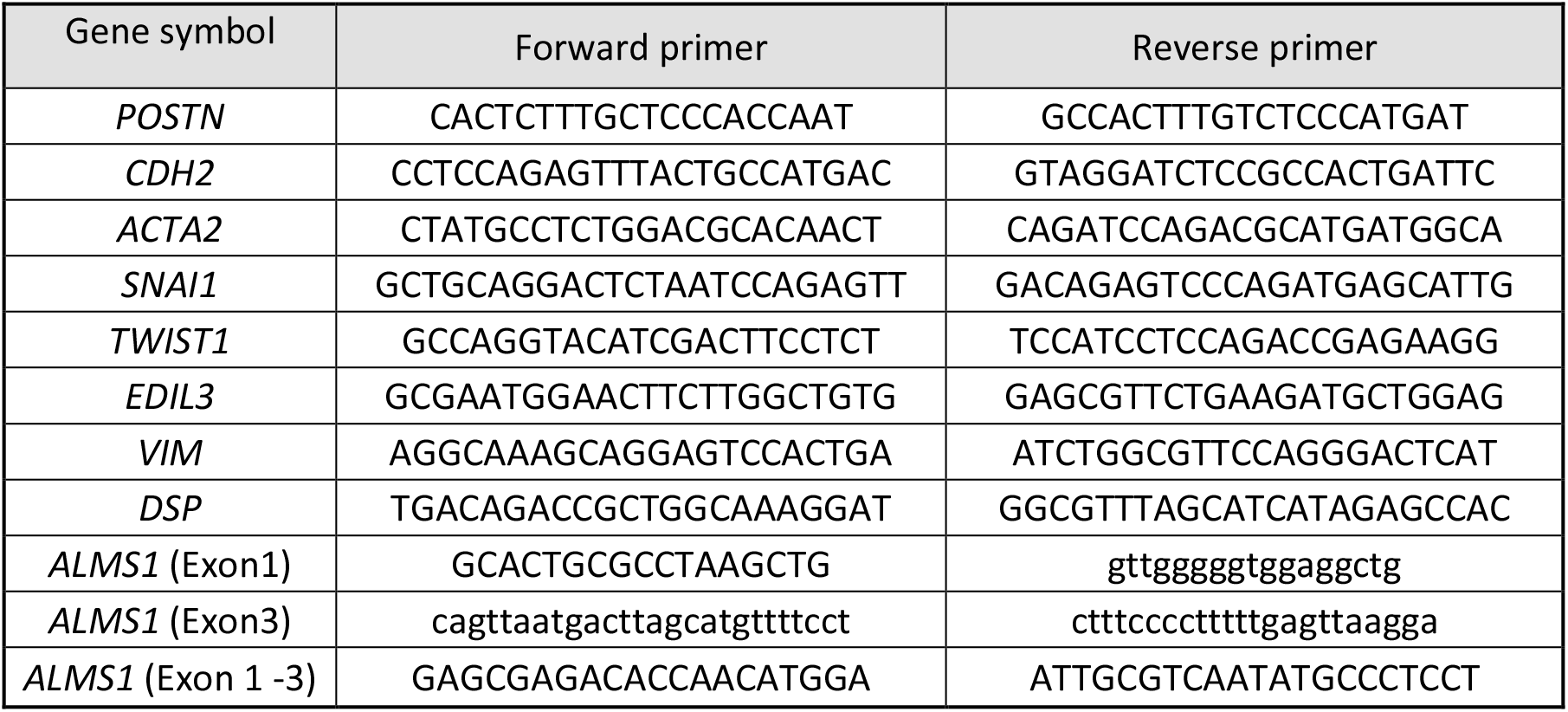
Sequences of the primers used for the evaluation of qPCR expression of epithelial-mesenchymal transition markers and genotyping of the Knockout lines.

### Wound healing assay

BJ-5ta and HeLa (WT and KO) were seeded in a 24-well plate (6 wells per cell line and genotype) at a concentration of 1×10^5^ and cultured until a 90-95% of confluence. Cells were serum-starvated (2%FBS for HeLa cells) for 24 hours. After that, rh-TGF-β1 (2 ng/mL) was added to half of the wells and a wound performed with a 200 μL pipette tip. Cells were incubated at 37°C 5% CO_2_ and images were taken at 0 and 24 hours with the microscope inverted Nikon TiU. Images were analysed with the imageJ program [28] and the plugin developed by Suarez-Armendo et al [29]. Finally, the percentage of closure was measured with respect to the wound area at time 0.

### SMADs activation assays

For p-SMADs/SMADs activation assays, BJ-5ta and HeLa (WT and KO) cells were plated in 100mm dishes (Corning, NY, USA) and incubated in DMEM:199 2P/S% without FBS and DMEM 2% P/S, 2%FBS for 24 hour, respectively. Then, TGF-β pathway were stimulated adding rhTGF-β1 ligand at 2 ng/mL (final concentration in dish) for 0, 10, 30, and 90 min in BJ-5ta and 0 and 30 min in HeLa. After stimulation, cells were washed 3 times with PBS, harvested using a scraper, and centrifuged in a Sigma^®^ 1–14 K at 4°C, 7,400×g for 10 min. Then, pellets were lysed on ice for 10 min in 200 μL of RIPA lysis buffer containing 1 mM sodium orthovanadate as phosphatase inhibitor (Sigma-Aldrich, Misuri, USA) and 0.1% (v/v) protease inhibitor cocktail (Merck, Darmstadt, Germany). Lysates were centrifuged at 4°C, 14,500×g for 30 min, and the supernatant was collected in 1.5mL low binding tubes (Thermo Fisher, Waltham, EE. UU). The remaining pellet was discarded. Quantification was performed using Bradford assay in microplates with Bio-Rad protein assay and samples were stored at −80°C until analysis.

### SDS-PAGE, Western Blotting, and Quantification

A total amount of 20μg from each sample were mixed with 6,25μL of Laemmli Buffer 4X (Biorad, Hércules, EE.UU), 1,25μL β-mercaptoethanol (BME) and a volume of H2Od up to a final volume of 25μL. Then, the mixes were boiled at 97°C for 5 min and loaded in a Mini-PROTEAN^®^ TGX™ Precast Gel (Biorad, Hércules, EE.UU) at 12% acrylamide/bis-acrylamide. The gel was migrated for 1 hour at 150V. Then, gels were transferred to a 0.2 μm PVDF membrane using the Trans-Blot Turbo transfer system (Biorad, Hércules, EE.UU) following the mixed molecular weight (MW) protocol. After transfer, membranes were blocked with in-house tris-buffered saline, 0.1% Tween20 and 5% milk (TBS-T 5%) for 90 min. Incubation with the primary antibody was kept at 4°C overnight. The primary antibodies used in this assay were: anti-SMAD3 (1:1000, Abcam, 40854), anti-p-SMAD3 (1:1000, Abcam, 63403), anti-SMAD2 (1:1000, Abcam, 40855), anti p-SMAD2 (1:1000, Cell Signaling Technology, 3101).

Next day, incubation with secondary antibody was performed by 45min in TBST 5%, after washed the membrane 3 times in TBST 5%. The secondary antibody used in this assay was goat anti-rabbit IgG H&L (HRP) (Abcam, 205718) at 1:5000 for p-SMAD2 and p-SMAD3 and 1:15000 for SMAD2 and SMAD3. Finally, for protein visualization Clarity western ECL substrate (Biorad, Hércules, EE.UU) was used and membranes were exposed in the ChemiDoc system (Biorad, Hércules, EE.UU). Quantification of protein bands was carried on Image Lab software (Biorad, Hércules, EE.UU) using the total protein standardisation method.

Western Blotting membranes were stripped and re-probed using Restore ™ PLUS Western Blot Stripping Buffer (Thermo Fisher, Waltham, EE. UU) according to the manufacturer’s protocol. Incubation for 15 min in the buffer was followed by 3 washes with 1X TBS. Then, the membranes were re-blocked and re-labelled following the previously described protocol.

### Statistical analysis

On one hand, mitochondrial activity, apoptosis, cell cycle, qPCR, Western Blot and wound healing data visualization and statistical analysis were performed with the software GraphPad Prism 8.00.

Mitochondrial activity was analysed using a linear regression model. Comparison of fluorescence increase with cell concentration did not show significant differences between both slopes (α = 0.05). The apoptosis experiment results were analysed with 2ways-Anova to compare each different population between genotypes. A Sidak’s multiple comparisons test was applied to correct the analysis. Finally, cell cycle results were analysed with a two-tailed unpaired t-test for each population (G2/M, S and G0/G1).

On the other hand, proteomic data analysis included different statistical analyses and data transformations along the pipeline. Initially, the data were normalised by variance stabilizing transformation (VSN). The data imputation for missing values was performed like missing not at random (MNAR) using the Minprob method with FDR < 0.01. Finally, differential expression analysis was performed using protein-wise linear models combined with empirical Bayes statistics implemented in the limma package, FDR < 0.05.

A third analysis was performed for the qPCR data. After normalization, fold changes (FC) were calculated concerning the geometric mean of the expression of each gene in the biological triplicates for *ALMS1* +/+ at 24h. Significant differences were calculates using a multiple t-test with a significant threshold of FDR < 0.05.

Finally western-blot data were normalized using total protein method following the methodology used by Clement et al. [22], [30] and as modified by Gürtler et al. [31]. After normalization FCs were calculated respect WT at 30 min of TGF-β1 stimulation. Significant differences were calculates using 2-way-ANOVA with a significant threshold of adj. p-value (FDR) < 0.05. For wound healing assay, a 2-way-ANOVA was applied and a significant threshold of adj. p-value (FDR) <0.05 was set.

## RESULTS

### *ALMS1* depletion does not alter mitochondrial basal activity despite generating G2/M cell cycle arrest in HeLa cells

*ALMS1* is a structural centromere protein [5] that controls the structure of the cytoskeleton [32]. Since *ALMS1* depletion causes cell cycle elongation [8], [33] it is expected that mitochondrial activity will also be altered, as both processes are dependent on each other and regulated by cytoskeletal dynamics. [34].

We evaluated the different phases of the cell cycle using the fox synchronous algorithm [35] to determine the Gaussian distribution of the different populations. DAPI fluorescence (461 nm) and events count (cells) were displayed in the plots (Figure 1.A). After 24h of incubation, HeLa cells ALMS1 −/− shown a significantly reduction (p-value < 0.05) in S phase from 24.26 % ± 1.32 in *ALMS1* +/+ to 19.27 % ± 1.69 in *ALMS1* −/−. The cell cycle arrest was caused by a significant increase in the G2/M population (p-value < 0.01), from 1.6%± 0.62 in *ALMS1* +/+ to 4.88 % ± 0.80 (Figure 1.B). The results in the G0/G1 population did not show significant differences between genotypes. Despite this, the basal mitochondrial activity did not show a significant difference between *ALMS1* +/+ and *ALMS1* −/− (Figure 1.C).

**Figure 1.**
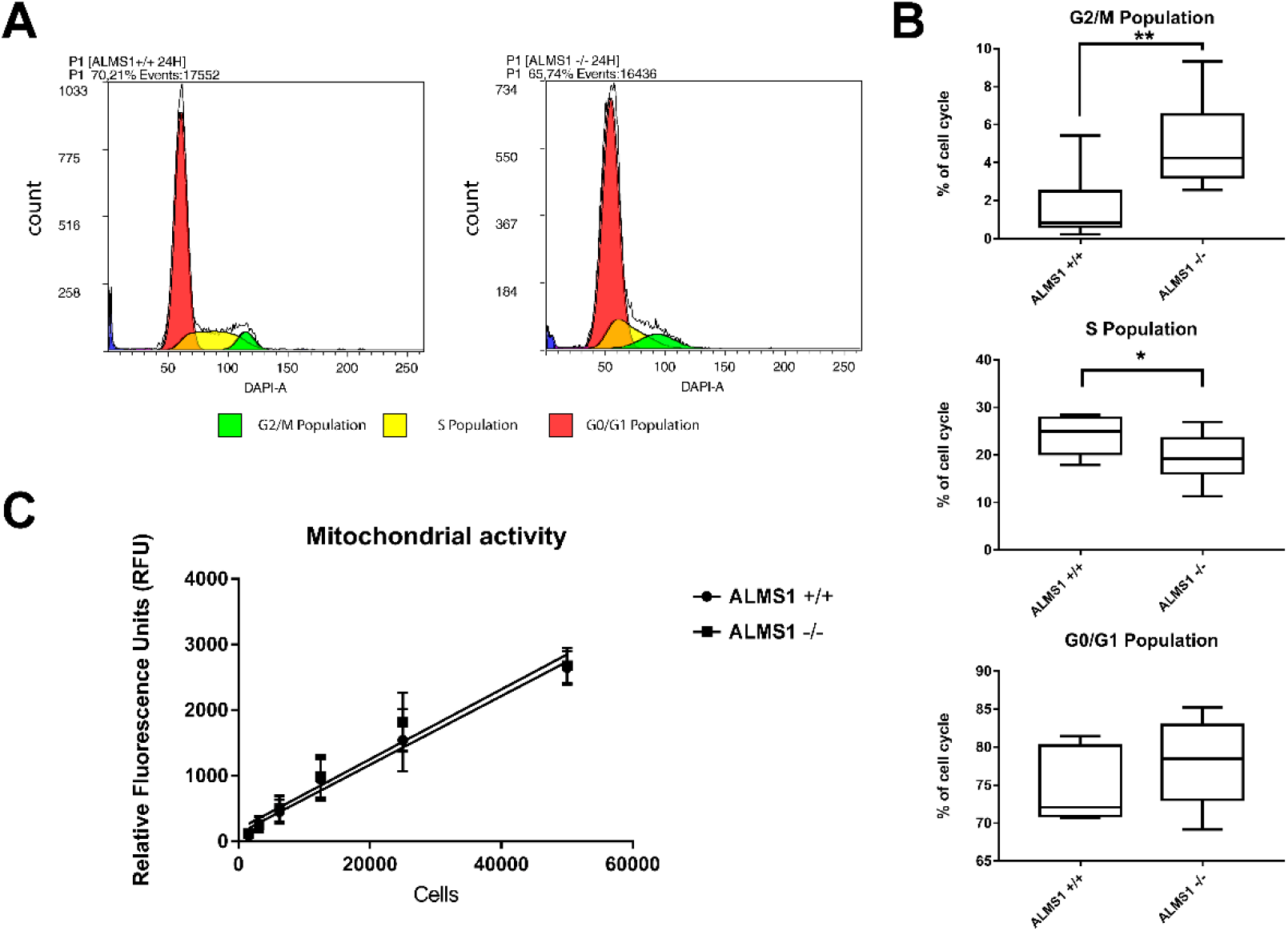
Cell cycle assay and mitochondrial activity results. **(A)** Cytometry plots with DAPI fluorescence and events count. Colors correspond with each population defined by fox synchronous algorithm. **(B)** Box-plot for each population previously defined in the cytometry plot that represents the percentage of cell cycle populations from parent population. **(C)** Linear regression between relative fluorescence units (RFU) and cell concentration in both genotypes *(ALMS1 +/+* and *ALMS1* −/−). This experiment was carried out in triplicate and repeated 3 times on different days (N=3, n=3). *: p-value < 0.05, **: p-value < 0.01.

### The lack of *ALMS1* in HeLa prevents apoptosis induced by C2-ceramides and Thapsigargin

Accumulation of cells in G2/M phase triggers apoptosis by different pathways such as caspases [36], [37]. We evaluated whether these cells were sensitive to apoptosis in response to an ER stress-inducing stimulus (Thapsigargin; THAP) [38], and a pro-apoptotic sphingolipid such as C2-ceramide.

As in the cell cycle assay, initial isolation of the cell population (P1) of interest was performed. Plots were represented with the FL4 (7-AAD) fluorescence in the y-axis and FL1 (Annexin V) fluorescence in the x-axis, both on a logarithmic scale. Thresholds for different populations (viable, early apoptotic, late apoptotic, and necrotic) were defined by a previous compensation analysis using THAP-like positive control of cell death. Finally, experiments were performed by triplicates in the plate (n=3) and in 3 alternative days (N=3).

Genotypes *ALMS1* +/+ and *ALMS1* −/− showed significant differences in viable (44.02% and 69.49%, respectively) and early apoptotic (43,47% and 23,44%) populations in cells stimulated with C2-C (Figure 2.A-B). Moreover, cells stimulated with THAP also showed significant differences between genotypes in viable (49,56% and 71,33%), early apoptotic (18,25% and 8,56%) and late apoptotic (25,09% and 11,93%) populations (Figure 2.C-D). These results indicate a resistance to apoptosis for these stimuli due to the lack of the *ALMS1*.

**Figure 2.**
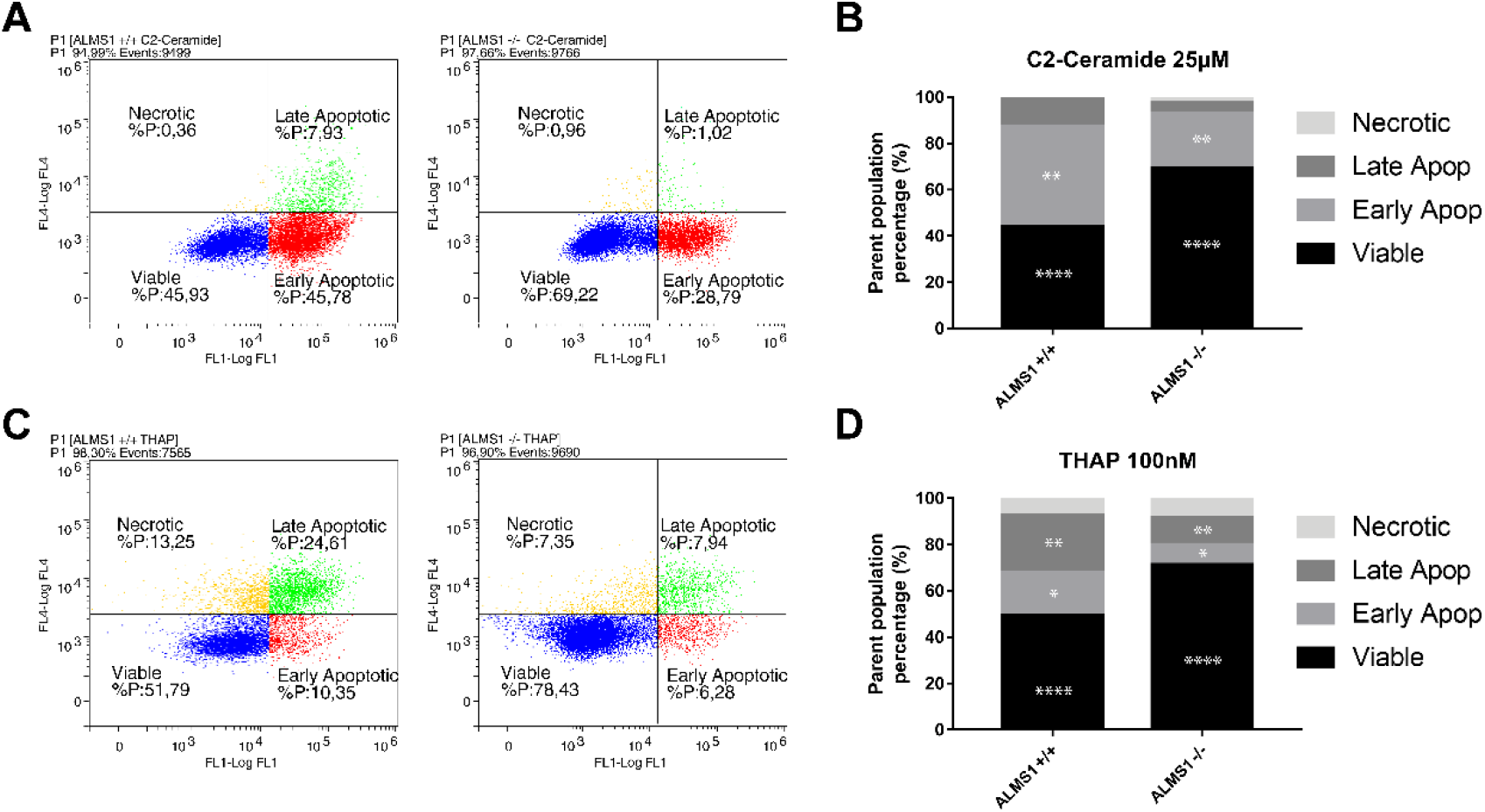
Apoptosis assay by annexin-V IP. **(A)** Cytometry plots for C2-ceramide (C2-C) with Log FL4 channel (IP) vs Log FL1 channel (Annexin-v). **(B)** Stacked-bar graph for cells treated with C2-C in a concentration of 25 μM. Percentages of each population are represented from a parent population previously defined. **(C)** Cytometry plots for thapsigargin (THAP) with Log FL4 channel (IP) vs Log FL1 channel (Annexin-v). **(D)** Stacked-bar graph for cells treated with THAP in a concentration of 100nM. Percentages of each population are represented from a parent population previously defined. This experiment was carried out in triplicate and repeated 3 times on different days (N=3, n=3). *: adj. p-value < 0.05, **: adj. p-value < 0.01, ***: adj. p-value < 0.001, ****: adj. p-value < 0.0001.

### The proteomic profile in HeLa cells reveals *ALMS1* like a regulator of cell adhesion through the TGF-β pathway

TGF-β is a pathway that plays a key role in the control of cell proliferation and apoptotic processes [14]. Since signal transduction through this pathway occurs via the primary cilium [13], [39], the lack of *ALMS1* could explain the aberrant processes seen above.

We evaluated how *ALMS1* depletion affects the signal transduction through stimulated TGF-β pathway. In this bulk analysis, we obtained a total of 86 differentially expressed proteins (DEP) (Figure 3.A), 31 (36%) over-expressed and 55 (64%) under-expressed. The variation in protein expression in this dataset can be explained mainly by a single principal component (Figure 3B). This would make sense as the main difference between these two genotypes would be the presence or absence of the *ALMS1* gene. From this dataset, a total of 76 proteins were associated with genes (Figure 3.C).

**Figure 3.**
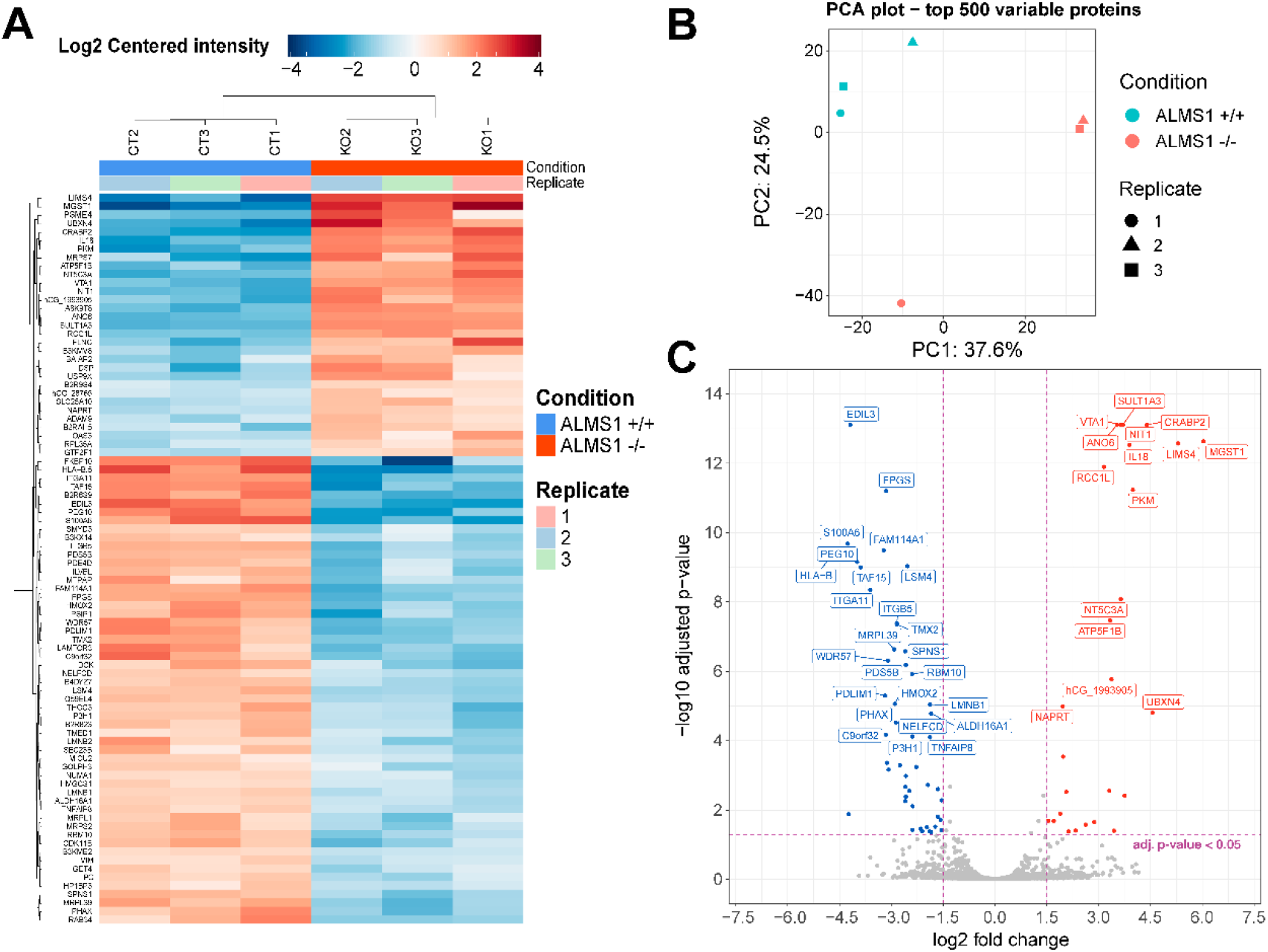
Differential expression analysis for proteomic data in HeLa. **(A)** Heatmap for all proteins (associated and not associated with genes) differentially expressed. Plot shows the expression pattern for the 3 biological replicates of genotype *ALMS1* +/+ (CT) and genotype *ALMS1* −/− (KO). **(B)** Principal components analysis (PCA) for the top 500 variable proteins with all biological replicates for each genotype. **(C)** Volcano-plot showing the top 40 most significant genes associated with proteins. Proteins with no clear gene association were removed from this plot. Significant thresholds: log2 FC > 1.5, adj p-value < 0.05. This experiment was carried out in triplicate (n=3).

Based on the DE of protein-coding genes (PCG) we performed an enrichment analysis obtaining a total of 9 GO terms significantly overrepresented (adj. p-value < 0.05) (Figure 4. A-B). The majority of these PCG were associated with focal adhesion, cell-cell or cell-substrate interactions. This result suggests the implication of *ALMS1* in the control of adhesion processes. In addition, we calculated the z-score of these GO terms (Figure 4.A), showed a significant decrease in process related to the ability of these cells to properly migrate or adhere to each other or their matrix.

**Figure 4.**
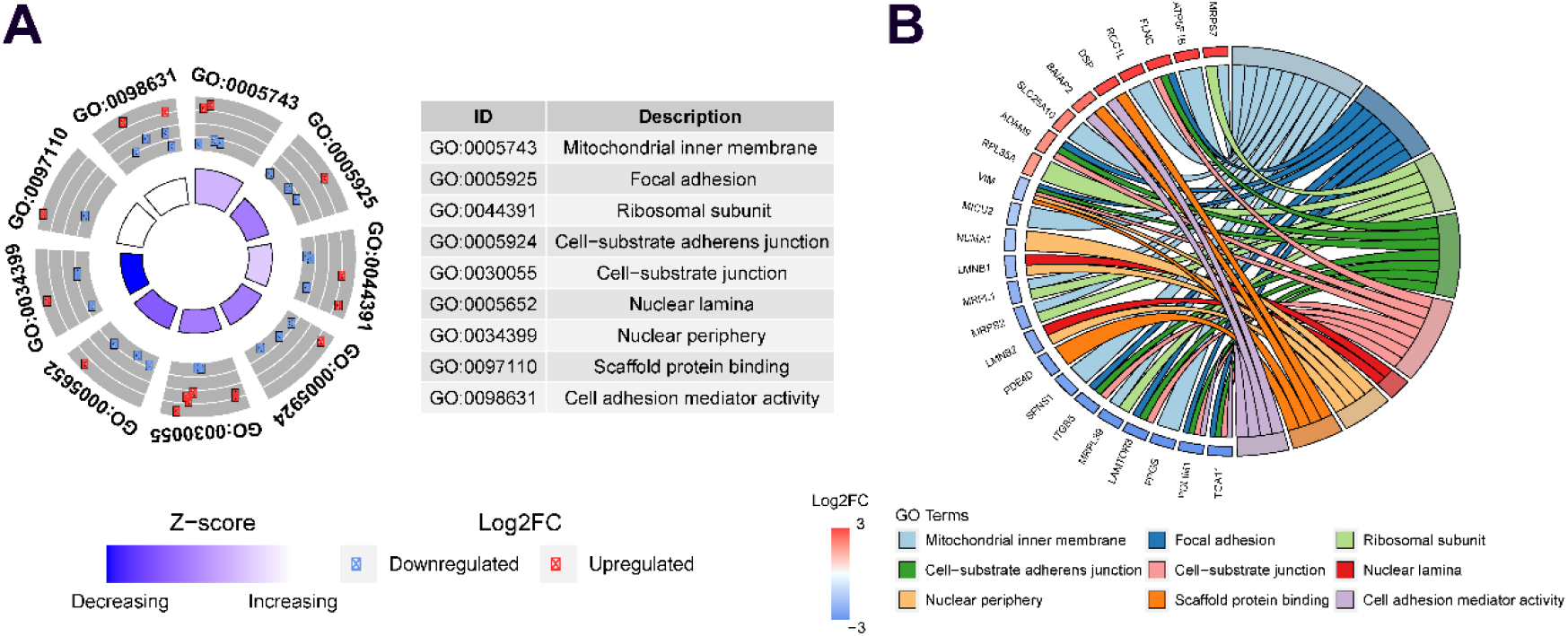
Enrichment analysis based on Go terms for proteomic data. **(A)** GoCircle-plot that includes significant over-represented GoTerms in a table (ID and Name) and scatter plot with FC of any gene related with each GoTerm. Moreover, this plot includes Z-score which is an estimation of up-regulation or down-regulation in the Goterm. **(B)** Circus-Plot applied to Goterms to relate any Gene name with the corresponding GoTerm name. The plot also shows the FC of each gene, and the list is sorted by FC from the most over-expressed (3) up to the most under-expressed (−3).

Moreover, genes such as vimentin *(VIM)* and desmoplakin *(DSP),* common biomarkers of EMT, were associated with these GO terms (Figure 4.B). Interestingly, these genes exhibited an expression pattern opposite to that observed when EMT process is activated.

Finally, *ITGB5* and *ITGA11,* two integrins whose activation plays an important role in the normal development of EMT, proliferation and invasion are inhibited [40], [41].

### *ALMS1* depletion alters the expression pattern of TGF-β1-induced EMT markers in HeLa

Proteomic analysis showed alterations in several adhesion processes coupled with aberrant expression of EMT-related genes *(VIM, DSP, ITGB5* and *ITGA11).* We measure the main EMT markers in our *ALMS1-*deficiency HeLa cells and compare it with another *ALMS1* model in the hTERT-BJ-5ta line.

We evaluated the expression of the principal EMT biomarkers *(SNAI1, CDH2, TWIST1, ACTA2, VIM,* and *DSP*) and other genes related to EMT *(POSTN, EDIL3).* We perform stimulations at different time points (24, 48 and 72 h for HeLa and 24 and 48 h for BJ-5ta) with rh-TGF-β1 (2ng/mL) stimulation.

In HeLa (Figure 5.A) at 24 hours of stimulation *POSTN*, *SNAI1*, *VIM*, *ACTA2* and *EDIL3* were found under-expressed, while *TWIST1* and *DSP* were over-expressed. This data confirms the expression pattern previously detected in the proteome analysis related to the genes *VIM, DSP* and *EDIL3* (Figure 5.B). At 48 hours of stimulation, the over-expressed genes *CDH2, TWIST1* and *DSP*, in addition to *EDIL3* and *VIM* inhibition, showed significant differences (Figure 5.A). No other tested gene were significant. Finally, at 72 hours of stimulation the expression pattern was like the previous stimulation time points. However, the data dispersion was too high, and no significant differences were detected.

**Figure 5.**
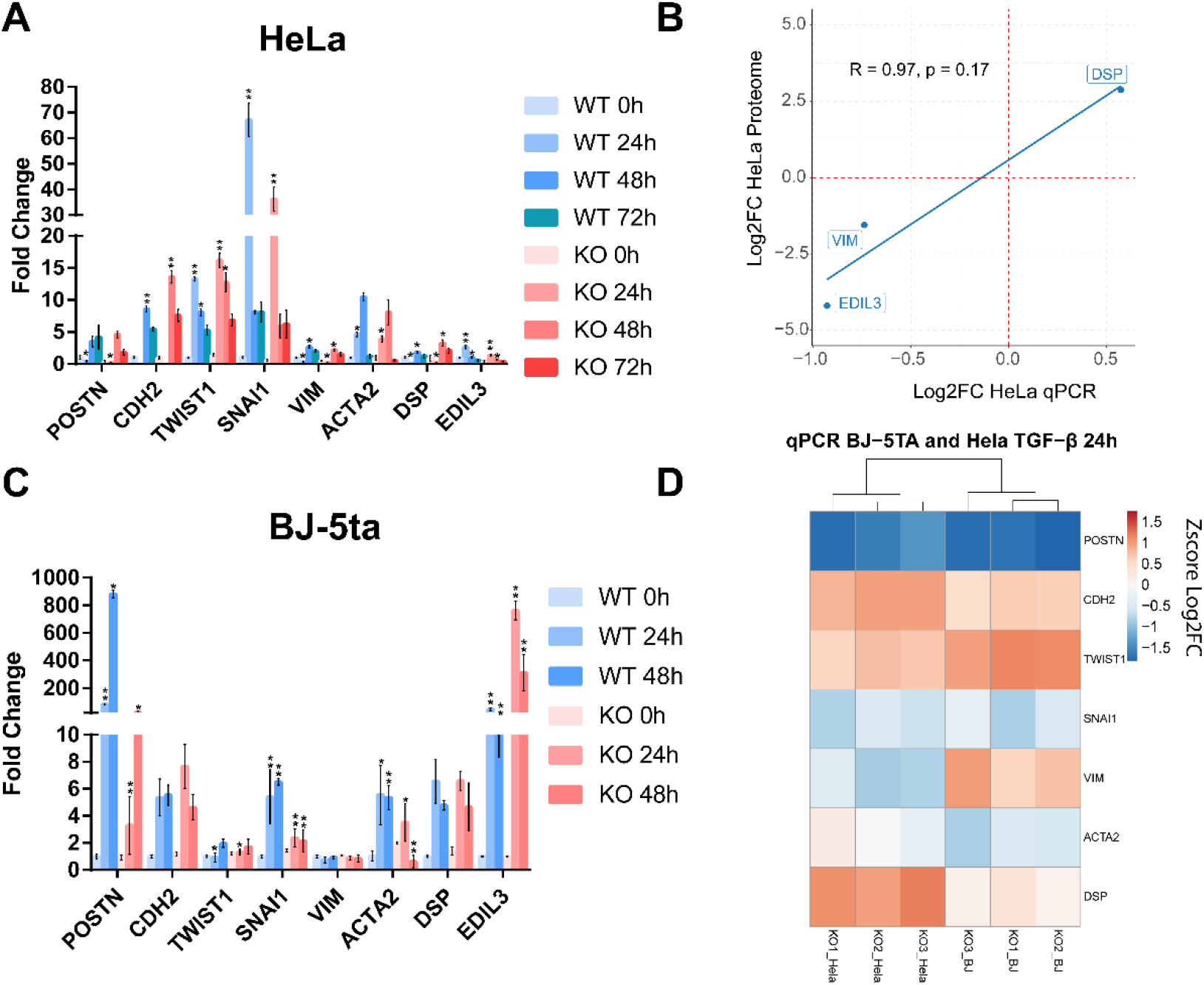
Expression measurement by qPCR of different EMT biomarkers with TGF-β stimulation at **(A)** 0, 24, 48, and 72 hours in HeLa. **(B)** Correlation plot between the expression of three EMT markers in the HeLa proteomic analysis and the HeLa qPCR assay. The expression is represented based on the log2 FC and the R coefficient using Pearson’s method. **(C)** Expression measurement of EMT markers by qPCR in BJ-5ta after 0, 24 and 48 hours of stimulation with TGF-β1 ligand **(D)** Comparative heatmap with Z-score values per sample of EMT markers in HeLa and BJ-5ta after 24 hours of TGF-β1 stimulation. These experiments were performed by triplicate (n=3). *: adj. p-value < 0.05, **: adj. p-value < 0.01.

In BJ-5ta (Figure 5.B) no inhibition of *VIM* or overexpression of *DSP* were detected. However, inhibition of *POSTN* and the transcription factor (TF) *SNAI1* was constant in both cell lines as well as the overexpression of *TWIST1. ACTA2* also showed decreased levels in HeLa but not as significant as in BJ-5ta. After Z score normalization of log2FC both cell lines showed a similar expression pattern of these markers (Figure 5C). EDIL3 is not shown in the heatmap as FC is too high, so it could mask the differences of the rest of the markers.

### Absence of *ALMS1* decreases migration capacity in HeLa and BJ-5ta and reduce SMAD3 phosphorylation in BJ-5ta

We evaluated the canonical activation of the TGF-β pathway. SMAD2 and SMAD3 phosphorilation levels were analysed at 4 different times (0, 10, 30 and 90 min) in BJ-5ta and 2 different times (0 and 30 min) in HeLa after TGF-β1 stimuli. The data shown that the lack of *ALMS1* inhibits SMAD3 (Ser213) activation after 30min of TGF-β1 ligand exposure (p-value <0.01) but not affects to SMAD2 phosphorylation in BJ-5ta (Figure 6 A-D). In HeLa cells no alterations in canonical pathway signalling of TGF-β were detected (Figure 6 E, F). The p-SMADs/SMADs ratios did not differ from the p-SMADs/total protein ratios, data not shown.

**Figure 6.**
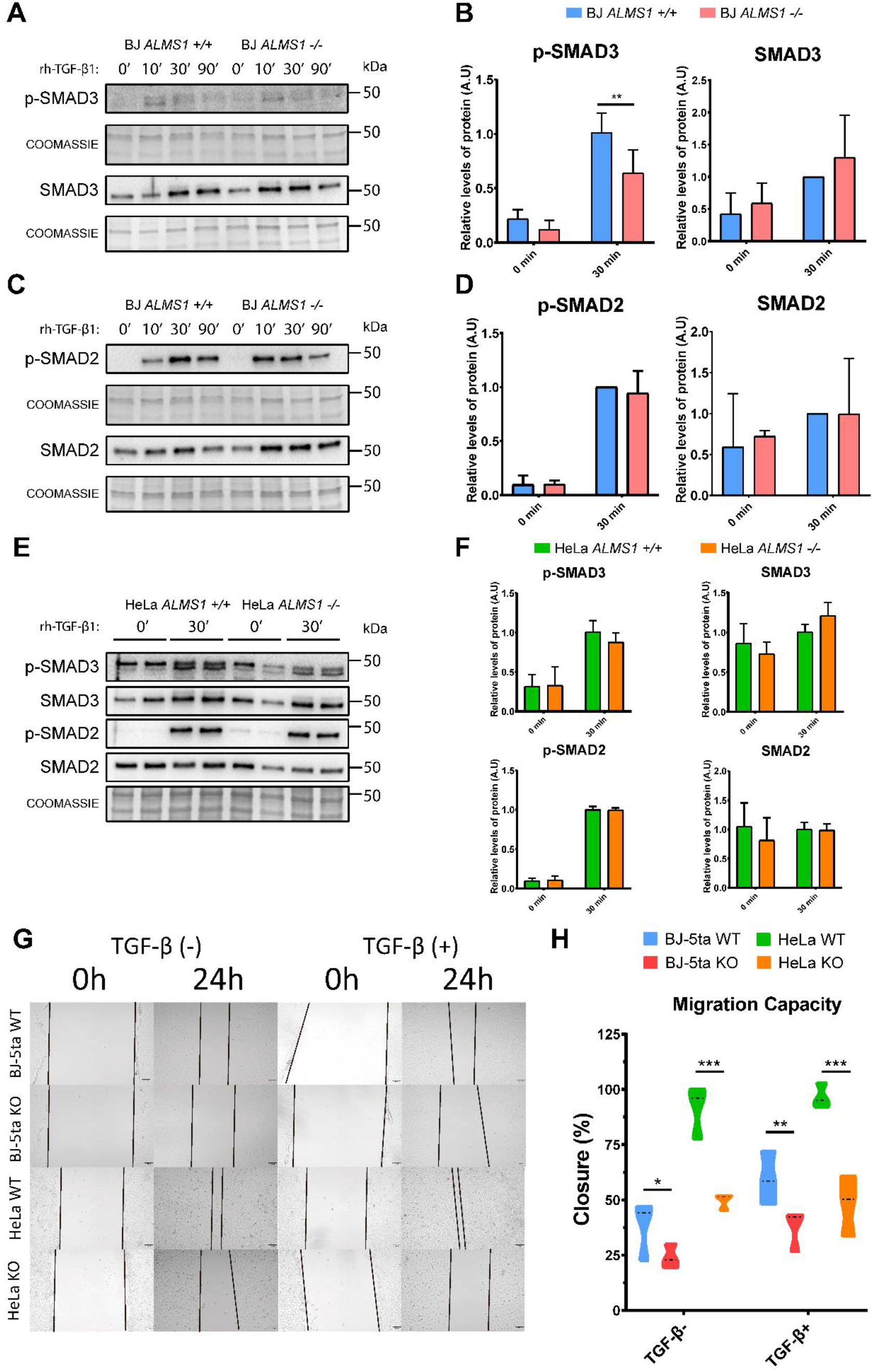
TGF-β canonical pathway activation assay and measurement of cell migration capacity in HeLa and BJ-5ta. **(A)** Representative western-blot of p-SMAD3 and SMAD3 levels after 0, 10, 30 and 90 min of stimulation with TGF-β1 (2 ng/μL) ligand and their corresponding loading controls (coomassie). **(B)** Quantification of band intensity in WB for p-SMAD3 and SMAD3 at time 0 and after 30 min stimulation with TGF-β1. (n=3). *: p-value < 0.05, **: p-value < 0.01 **(C)** Representative western-blot of p-SMAD2 and SMAD2 levels after 0, 10, 30 and 90 min of stimulation with TGF-β1 (2 ng/μL) ligand and their corresponding loading controls (coomassie). **(D)** Quantification of band intensity in WB for p-SMAD2 and SMAD2 at time 0 and after 30 min stimulation with TGF-β1. (n=3). *: p-value < 0.05, **: p-value < 0.01. **(E)** Representative western-blot of p-SMAD3, SMAD3, p-SMAD2 and SMAD2 levels after 0 and 30 min of stimulation with TGF-β1 (2 ng/μL) ligand and their corresponding loading control (coomassie). **(F)** Quantification of band intensity in WB for p-SMAD3, SMAD3, p-SMAD2 and SMAD2 at time 0 and after 30 min stimulation with TGF-β1. (n=2). *: p-value < 0.05, **: p-value < 0.01 **(E)** Representative images of wound healing assay in HeLa (WT and KO) and BJ-5ta (WT and KO) at 0 and24 hours in absence or presence of 2 ng/μL TGF-β1. **(F)** Quantification of the cell area of the image and how it decreased after 24 hours of incubation (n=3). *: p-value < 0.05, **: p-value < 0.01, ***: p-value < 0.001.

Finally, we measured cell migration capacity in both cell lines in the presence and absence of - rh-TGF-β1 ligand. (Figure 6 G, H). *ALMS1* depletion reduced migration capacity compared to the control line in both cell lines. This reduction is exacerbated after TGF-β stimulation in BJ-5ta while in HeLa the tendency is the same in the presence or absence of the stimulus.

## DISCUSSION

In this study, we obtain similar results as previously described for the ALMS1 deficiency such as cell cycle arrest [33] or resistance to apoptosis [8] in a new cell type. Moreover, we reported new insights to understand the role of *ALMS1* in cell function and TGF-β signalling Cell cycle arrest in the G2/M phase (Figure 1. A-B), also described in *ALMS1*-knockdown cardiomyocytes [33], seems to be a cell process triggered by *ALMS1* depletion. Several studies have already linked cell cycle arrest in the G2/M phase to the TGF-β pathway and the development of fibrosis [42]–[44].

Normally, cell cycle arrest in both the G2/M and G0/G1 phases is often accompanied by the activation of apoptotic processes [34], [45]–[47]. In contrast, depletion of *ALMS1* seems to generate apoptosis resistance against THAP and C2-C in HeLa cells (Figure 2), which is in accordance with that reported by Zulato et al., [8] in patient fibroblasts. THAP is a non-competitive inhibitor for Ca^2+^-ATPases located in the endoplasmic reticulum (SERCA). This, inhibits Ca ^2+^ reflux to the lumen of the endoplasmic reticulum, generating endoplasmic reticulum stress (ER stress) [48]. At the same time, ER stress activates a signalling network called Unfolded Protein Response (UPR) which alleviates this stress and promotes cell survival, restoring ER homeostasis [49]. At this point, *ALMS1* depletion could be acting at two levels: inhibiting THAP action (ER stress) or over-activating UPR and generating apoptosis resistance. Because of the role of *ALMS1* in the centrosome and its relation with endocytic trafficking [32], its depletion could probably alter the structure of the membrane, inhibiting the signalling transduction of ligands like THAP. THAP also induces apoptosis impairing cytoskeleton dynamics and disorganization of F-actin via mTORC1 inhibition in the mTORC1-RhoA-Limk-Cofilin-1 axis in A549 cells [50]. *ALMS1* depletion already causes disorganization of the F-actin cytoskeleton [32] but does not lead to cell death. Probably, the mTOR pathway through mTORC1 was inhibited in *ALMS1* −/− cells, but in this case, other mechanisms could prevent cells from reaching apoptotic stages. This would be supported by the inhibition of LAMTOR3 (Figure 4.B), a protein involved in the formation of the Ragulator complex that activates mTORC1 [51].

C2-C is a sphingolipid that induces caspases-dependent apoptosis, inhibiting PI3K/AKT pathway and increasing the depolarisation of the mitochondrial membrane [52]–[54]. The over-activation of the PI3K/AKT pathway could explain the apoptotic resistance in the case of the C2-C [55], [56]. This fact would agree with the previous proposed mTORC1 inhibition, since phosphorylation of AKT/PKB at s473 leads to inhibition of mTORC1 mediated by FoxO transcription factors [57], which could be related to F-actin filament disorganisation previously described in Alström fibroblasts [32].

In addition to the role of *ALMS1* in G2/M phase cell cycle arrest and resistance to THAP- and C2-C-mediated apoptosis, this study suggests that *ALMS1* depletion leads to inhibition of TGF-β signalling pathway. This could be affecting downstream processes such as cell migration and adhesion.

After TGF-β stimulation, the lack of *ALMS1* alters the expression of several genes related to focal adhesion and cell-substrate adherens junction (Figure 4 A-B), two processes required for TGF-β1-induced EMT [58], [59]. This fact could be conditioning the activation of EMT, preventing cells from moving into a mesenchymal state characterised by an increased migratory capacity. Previous studies have demonstrated the relationship between signalling through the primary cilium and the EMT focusing mainly on signalling through the WNT and SHH pathway in various organs such as epicardial tissue, kidney epithelial cells, and pancreatic β-cells [31]–[34]. However, signalling through the TGF-β pathway has not yet been extensively studied in this context.

Moreover, we have found interesting candidates in the HeLa proteome profiling that could explain or be related to the mentioned cellular disturbances (Figure 3C). *S100A6,* under-expressed, is a positive regulator of proliferation, migration and angiogenesis [60]–[62], and also a regulator of endothelial calcium signalling [63]. *PKM,* which was over-expressed in HeLa cells, is associated with apoptosis resistance [64]. *ITGA11,* another gene under-expressed in this cell line and related to fibrosis generation, is expressed due to activation of the TGF-β pathway [65]. Finally, two known EMT markers, *VIM* and *DSP*, showed a reverse pattern to that expected when this process is activated. *VIM* is a key player in the regulation of EMT and its deficiency leads to inhibition of this process [66]. On the other hand, *DSP* is a protein involved in cell-cell junctions mediated by desmosomes. It is therefore considered a typical epithelial marker whose decrease is related to the passage of the cell from an epithelial to a mesenchymal state [67]. This could be first evidence suggesting inhibition of the TGF-β1-induced EMT when *ALMS1* is missing.

One of the most relevant detected genes was *SNAI1,* which is inhibited in both cell lines (HeLa and BJ-5ta). *SNAI1* and *TWIST1* are transcription factors that repress E-cadherin *(CDH1)* expression and activate the expression of N-cadherins *(CDH2)* [68], [69]. E/N-cadherins switch is involved in the induction of EMT [70], [71] through the TGF-β pathway [72]. Inhibition of *SNAI1* could be affecting the normal EMT in these cells. The decreased expression of *SNAI1,* together with the inhibition of *ACTA2* and *POSTN,* also suggests that TGF-β1-induced EMT would be down-regulated. *SNAI1* is up-regulated by SMAD3, which is phosphorylated when the TGF-β pathway is activated and forms a complex with SMAD2 [17]. The phosphorylated SMAD2-SMAD3 complex translocates to the nucleus, activating the expression of several genes related to cell migration and EMT such as *SNAI1* [73]. Because we detected consistent inhibition in both cell models of *SNAI1,* we sought to determine whether SMAD3 and SMAD2 activation was also compromised in BJ-5ta and HeLa models. In BJ-5ta, we conclude that *ALMS1* depletion inhibits SMAD3 phosphorylation but not affect SMAD2 phosphorylation (Figure 6 A-D). This fact could lead to downstream inhibition of *SNAI1* and affect migration and adhesion capacity in this cell line (Figure 6 G, H). In HeLa, no alterations in the activation of the canonical TGF-β pathway were detected (Figure 6 E, F). In addition, a decrease in cell migration capacity was observed, but independently of TGF-β, in contrast to BJ-5ta (Figure 6 G, H). This could explain that the involvement of ALMS1 in cell migration is a conserved process, but its regulation is a cell type-dependent process.

We have previously described that *ALMS1* depletion by siRNA inhibits SMAD2 phosphorylation in the hTERT RPE-1 cell line [10] but this was not detected in any of our KO models. The only alteration detected in the canonical pathway was the inhibition of SMAD3 activation in BJ-5ta. This difference could be due to the cell type or to differences between the KO and siRNA models.

This facts leads us to hypothesise that inhibition of p-SMAD3 could prevent the formation of the SMAD2/3 complex, affecting the expression of *SNAI1* in BJ-5ta. [20]. Whereas in HeLa, inhibition of *SNAI1* and cell migration capacity appears to involve other regulatory mechanisms.

## CONCLUSION

*ALMS1* appears to have a role in the regulation of cell cycle and apoptotic processes but does not affect mitochondrial reduction capacity in HeLa. *ALMS1* depletion affects signal transduction through the TGF-β pathway, altering the expression of several EMT-related genes in BJ-5ta and HeLa. However, the canonical (SMAD-dependent) pathway did not show major differences. Finally downstream processes such as cell migration and adhesion capacity appear to be compromised in the absence of *ALMS1.*

## Supporting information

Supplementary Table S1 gRNAs

Supplementary Table S2 Primers EMT markers

## Author Contributions

BB-M and DV designed the study. BB-M performed the experiments and analysed the data. EN-G assisted in performing apoptosis, cell cycle and mitochondrial activity assays. AI-R assisted in the quantification of the expression of EMT markers in the BJ-5ta model and in the cell migration assay. BB-M and DV wrote the manuscript. All authors have read the draft and provided approval for publication.

## Data Availability Statement

Proteomic data are available via ProteomeXchange with identifier PXD024964.

## Acknowledgments

We sincerely thank the Proteomics and Genomics services from Centro de Apoyo Científico-Tecnológico a la Investigación (CACTI) of University of Vigo and its specialists Paula Álvarez Chaver, Ángel Sebastián Comesaña, Verónica Outeiriño and Manuel Marcos for their guidance and advise. We also thank Mercedes Peleteiro Olmedo from Centro de Investigacións Biomédicas (CINBIO) from University of Vigo for the flow cytometry service.

## Supplementary Material and Methods

### CRISPR plasmids construction

Single guides RNAs (sgRNAs) for BJ-5ta model were designing using benchling to generate a deletion of 113pb in the exon 3 of *ALMS1* (ENSG00000116127). The 2 sgRNA (5’-CCTAGTCTGGGATGTATCCA – 3’ and 5’-GAACAGAAGAGATCTCTGTT – 3’) result from benchling were cloned into the plasmid lentiCRISPR v2 [1] following a modified protocol from that described by Shalem et al [2]. LentiCRISPR v2 was a gift from Feng Zhang (Addgene plasmid #52961). Initially, 12.5μL (5μg) of LentiCRISPR V2 were digested with 3μL Esp3I (BsmBI), 3μL FastAP and 6μL 10X FastDigest Buffer from Thermo Fisher. Final volume was set at 60μL. Plasmid digestion was checked by migration in an agarose gel at 1%.

sgRNAs were phosphorylated and annealed with its correspondent oligo mixing as follow: 1μL sgRNA (100μM), 1μL of its complement oligo, 2μL of 5X T4 Buffer (Canvax, Cordoba, Spain), 0.5 T4 PNK (NEB, Ipswich, EE.UU) and 5.5 μL distilled H_2_0 (H_2_Od). Mix reaction was incubated at 37ºC for 30min and 95ºC for 5min. The samples were then left to stand at room temperature for 2 hours and diluted at 1:50, 1:100 and 1:200 ratios.

The unpurified digested plasmid was ligated with the diluted sgRNAs as follows: 0.5μL digested LentiCRISPR v2, 2μL diluted sgRNA, 1μL T4 DNA Ligase (Canvax, Cordoba, Spain), 4.5μL H_2_Od. The mix was incubated at 37ºC by 30min.

Finally, Stable Competent E.coli (NEB, Ipswich, EE.UU) were transformed by thermal shock according to the manufactured protocol and incubated at 30°C overnight. After that, construct validation was carried out using forward primer for U6 promotor (5’-GACTATCATATGCTTACCGT-3’) and reverse sgRNA.

### Plasmids Transfection and Lentivirus production

293T were seeded in a concentration of 3×10^5^ into a 6-well plate and incubated overnight. Then, 293T cells were co-transfected with each CRISPR construct and the plasmids psPAX2 and pCMV-VSV-G [3] using lipofectamine 3000 (Thermo Fisher, Waltham, EE. UU). pCMV-VSV-G was a gift from Bob Weinberg (Addgene plasmid #8454) and psPAX2 was a gift from Didier Trono (Addgene plasmid #12260)

Initially, 7.5μL of lipofectamine 3000 were diluted in 125μL of Opti-MEM medium (Thermo Fisher, Waltham, EE. UU) by duplicate. Then a total amount of 5μg of ADN (1.6μL for each plasmid) were diluted at 250μL Opti-MEM. After that, diluted DNA and diluted lipofectamine were mixed in a 1:1 ratio, centrifugated at 1200rpm and incubated for 20min at room temperature. Finally, the 250μL of mix were applied to the wells containing the cells.

The cells were incubated with the mixture for 24 hours. Then, the mixture was removed and the cell was cultured for a further 72 hours for lentivirus production.

### Lentiviral transduction

After 72 hours of culturing 293T cells transfected for lentivirus production, the culture medium was collected in 15 mL tubes and filtered through a 45μm filter (Sartorious,). In addition, a solution of complete DMEM medium without P/S with polybrene was prepared at a dilution of 1:1000 (8μg/ml).

Then, 300μL lentiviral preparation were mixed with 200μL complete DMEM mídium with polybrene for each well. An additional 1 ml of DMEM with polybrene was added and the plates were centrifuged at 800g for 30 minutes. This step was repeated one more time. After the second centrifugation the mix with lentivirus was added again and the plates were incubated for 24 hours. Finally, mix was removed and cells were allowed to settle for 48 hours in DMEM mídium. They were then selected with puromycin (1ng/mL) for 5 days.

### Edition validation

After puromycin selection, clonal isolation and expansion we checked the lack of *ALMS1* gene by qPCR in StepOnePlus (Thermo Fisher, Waltham, EE. UU). Finally, we found that these cells had the expected exon 3 deletion by PCR with resolution of samples on a 2% agarose gel and followed by sanger sequencing.

## Supplementary figures

**Supplementary Figure S1.**
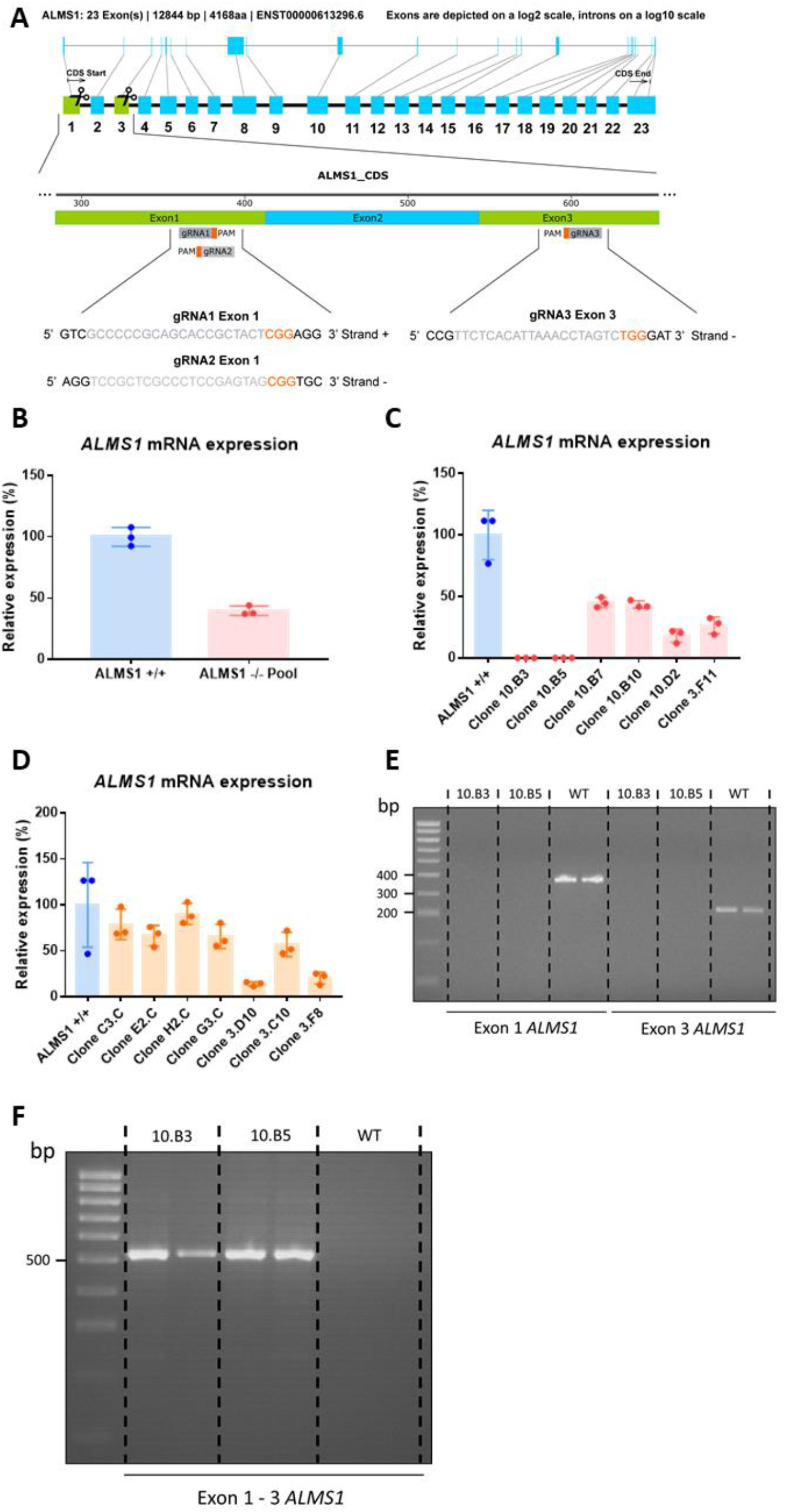
Characterisation of *ALMS1* −/− cell line HeLa. **(A)** Design of the model. **(B)** Relative gene expression of *ALMS1 +/+* and polyclonal cell line for *ALMS1 −/−* genotype. **(C)** Relative gene expression of ALMS1 +/+ and a first batch of clones for *ALMS1 −/−* genotype. **(D)** Relative gene expression of ALMS1 +/+ and second batch of clones for ALMS1 −/− genotype. **(E)** Strategy for amplification of exons 1 and 3 from genomic DNA in two clones of the *ALMS1* −/− (KO) line and the *ALMS1* +/+ control (WT). **(F)** Amplification strategy of the resulting sequence between exon 1 and 3 after removal of the sequencing between the 2 outermost gRNAs.

**Supplementary Figure S2.**
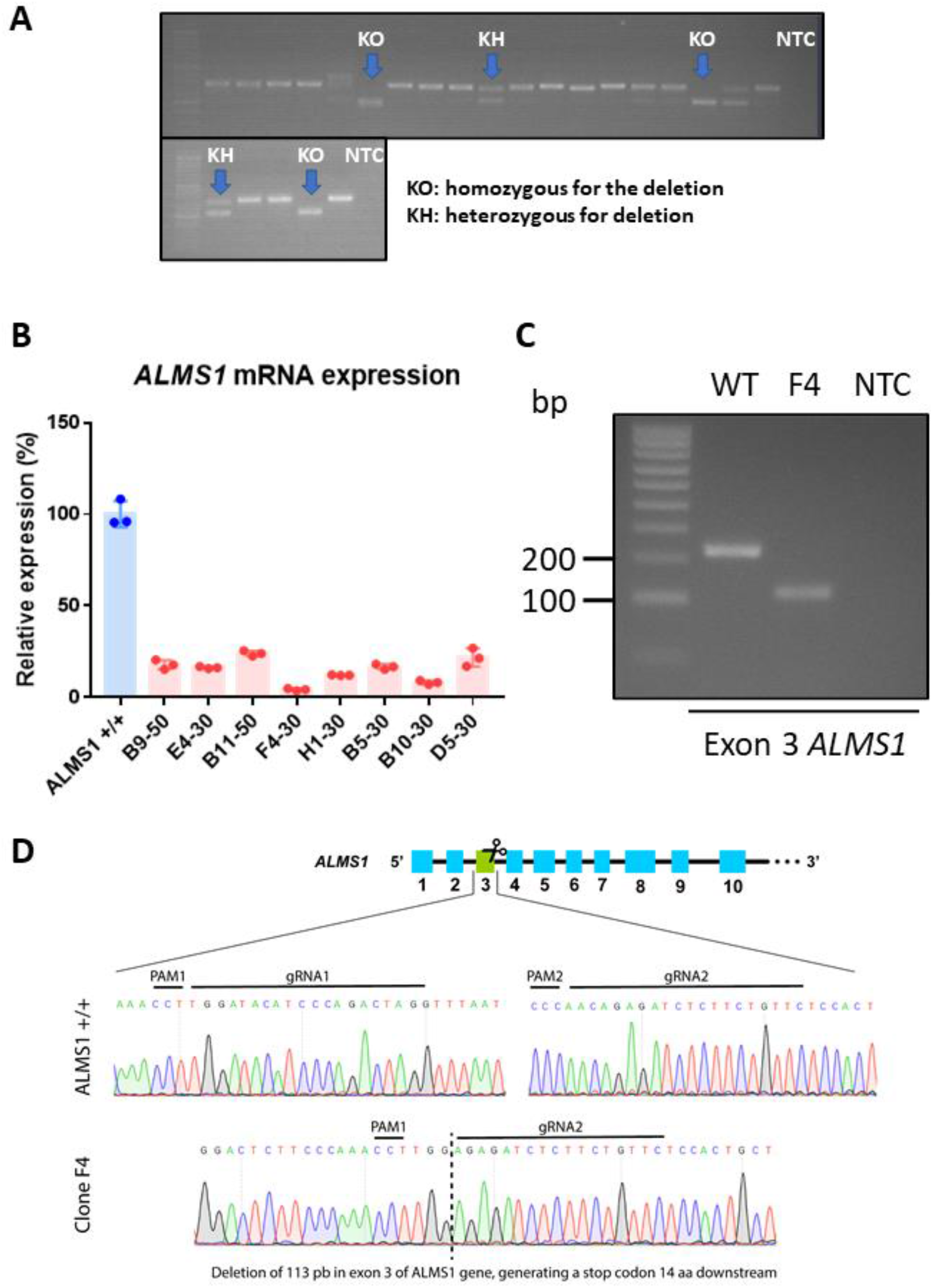
Genetic characterisation of *ALMS1* −/− cell line BJ-5ta. **(A)** Clone screening. **(B)** Relative gene expression of ALMS1 +/+ and a batch of clones for ALMS1 −/− genotype. **(C)** Agarose gel (2%) showing the exon 3 amplicon of the control line (ALMS1 +/+) and the F4 clone (ALMS1 −/−) which showed the lowest gene expression. **(D)** Characterisation of the F4 clone mutation by sanger sequencing.

